# The soil microbial food web revisited with metatranscriptomics - predatory *Myxobacteria* as keystone taxon?

**DOI:** 10.1101/373365

**Authors:** Sebastian Petters, Andrea Söllinger, Mia Maria Bengtsson, Tim Urich

## Abstract

Trophic interactions in the microbial food web of soils are crucial for nutrient and carbon cycling. Traditionally, protozoa are considered the major micropredators of bacteria in soil. However, some prokaryotes, such as *Myxobacteria* and *Bdellovibrio* are also famous for bacterivorous life style. Until recently, it was impossible to assess the abundance of pro- and eukaryotic micropredators in soils simultaneously. Using a metatranscriptomic three-domain profiling of small subunit ribosomal RNA we investigated the abundance of bacterivores in 28 datasets from eleven European mineral and organic soils of different climatic zones. In all soils, *Myxobacteria* comprised a significant proportion from 4 – 19% of prokaryotic 16S rRNA transcripts and more than 60% of all bacterivores in most soils. *Haliangiaceae* and *Polyangiaceae* were most abundant, while the name-giving *Myxococcaceae* were barely present. Other bacterial predators like *Bdellovibrio* were low abundant. Also Protozoan micropredator 18S rRNA transcripts, e.g. from *Cercozoa, Amoebozoa and Ciliophora*, were on average less abundant, especially in mineral soils. *Nematodes* were even less abundant. In addition, we applied a longitudinal approach to identify bacterivores during beech litter colonisation. Here, *Myxobacteria* showed prey-dependent, protozoa-like community dynamics during colonisation. Thus, their broad prey range and high abundance suggests a major influence of *Myxobacteria* on structuring the prokaryotic community composition in soil, and might warrant their classification as keystone taxon. Our results suggest the presence of an ecologically important “bacterial loop” in soil food webs, independent of protozoa and nematodes.

## Introduction

Predation of predators on prey is a key process in structuring community composition in ecosystems and in maintaining high biodiversity. Predator - prey interactions and dynamics among animals and consequences for ecosystem functioning have been studied extensively since the early days of ecology. While less visible and thus less acknowledged, predation is not foreign to the microbial world. Eukaryotic as well as prokaryotic microorganisms are known to prey on other microorganisms in marine, aquatic and terrestrial habitats as part of the microbial food web (Clarholm, 1985; Azam et al., 1983).

Protozoa are traditionally considered the main microbial predators of bacteria, a notion that stems from the fact that, unlike in bacteria, where it is somewhat “exotic”, predation is a common lifestyle among protozoa. Predatory protozoa are known from both aquatic and soil environments and have been considered a key-component of the “microbial loop” responsible for the remineralisation of nutrients (Bonkowski, 2004; Clarholm, 1985). Whereas protozoa in the aquatic system have been well characterised, both in terms of their identity and population size, research in soils has been much more hampered, since no adequate molecular tools have been available for a long time, cultivation is often difficult, and direct microscopic observations are impossible (Geisen et al., 2015).

Much fewer prokaryotic species are considered predatory, although a predatory lifestyle in prokaryotes probably evolved prior to its development in eukaryotes. Several bacterial predators have been identified, with more and more taxa exhibiting a predatory lifestyle being recognized recently. These include *Myxobacteria, Lysobacter, Bdellovibrio* and like organisms (BLO), *Vampirococcus,* and *Dapterobacter*, among others (Reichenbach, 1999). Especially the *Myxobacteria*, with their ‘wolf pack hunting’ strategy, are known micropredators since more than 70 years ago and have been isolated from soils world-wide (Keane & Berleman, 2016; Reichenbach, 1999).

It has been until recently impossible to assess bacterial and protist community composition with the same methodology. Although PCR amplicon approaches enabled the study of both groups separately, a direct comparison of their relative abundances was not possible due to the absence of universal primers that would tackle all groups without bias. However, these obstacles are avoided when applying random hexamer-primed reverse transcription as in metatranscriptomics approaches that target SSU rRNA of organisms from all three domains of life (Urich et al., 2008). Furthermore, these rRNA transcripts are indicative of ribosomes and thus are likely derived from metabolically active cells and can be considered markers for living biomass. The generated cDNA fragments originate from different regions of the SSU rRNA molecule unlike PCR primed specific sites, and are therefore insensitive to the presence of introns or primer mismatches, when PCR primers are applied.

We have recently used this PCR-free metatranscriptomics approach to reveal the diversity of the active soil protist communities within five different natural soil systems in Europe, including forest, grassland and peat soils as well as beech litter (Geisen et al., 2015).

Here we have focused on other groups of microbial predators - predatory bacteria. We have assessed the relative abundance of SSU rRNAs from bacterial groups known to exhibit a predatory lifestyle in these soils. Metatranscriptomics enabled the direct comparison of SSU rRNA transcripts from bacterial and protozoan micropredators and revealed that potentially predatory bacteria, especially *Myxobacteria*, were abundantly detected in all soils, while protozoa abundances were much more variable. The underlying causes and consequences for our perception of microbial predation in soils are discussed and an alternative model of the soil microbial loop is put forward.

## Material and Methods

### Data acquisition

The investigated metatranscriptomes had been obtained from different previous studies on a range of European soils (Table 1). These included 4 samples from organic peatland, 3 samples from organic floodplain, 3 samples from gleic fluvisol, 3 samples from mineral grassland, 2 samples from organic forest litter, 4 samples from mineral forest soil, and 3 samples each from 3 different mineral shrubland soils. RNA, cDNA and sequences were obtained as previouly described (Beulig et al., 2016; Epelde et al., 2015; Geisen et al., 2015; Tveit et al., 2013; Urich et al., 2008).

**Table 1.** Context data for relevant sampling sites (changed from Choma *et al.*, 2016)

| Site | Peatland soil<br>"Knudsenheia" | Peatland soil<br>"Solvatn" | Mofette | Mofette<br>reference | Rothamsted<br>grassland | Rotbühl | Forest Litter | Forest Soil | Mine L | Mine M | Mine H |
| --- | --- | --- | --- | --- | --- | --- | --- | --- | --- | --- | --- |
| Abbreviation | PsK | PsS | MO | MR | RS | RB | FL | FS | MiL | MiM | MiH |
| Location | Ny-Ålesund,<br>Norway<br>(Svalbard) | Ny-Ålesund,<br>Norway<br>(Svalbard) | Hartoušov,<br>Czech<br>Republic | Hartoušov,<br>Czech<br>Republic | Rothamsted,<br>United<br>Kingdom | Darmstadt,<br>Germany | Vienna woods,<br>Austria | Vienna woods,<br>Austria | Coto Txomin,<br>Spain | Coto Txomin,<br>Spain | Coto Txomin,<br>Spain |
| Climatic zone | Arctic | Arctic | Temperate | Temperate | Temperate | Temperate | Temperate | Temperate | Temperate | Temperate | Temperate |
| Biome | Fen wet land | Fen wet land | Floodplain | Floodplain | Grassland | Grassland | Temperate<br>deciduous<br>forest | Temperate<br>deciduous<br>forest | Shrubland | Shrubland | Shrubland |
| Dominant vegetation | Mosses | Mosses | *Filipendula*<br>*ulmaria* | *Deschampsia*<br>*cespitosa,*<br>*Eriophorum*<br>*vaginatum* | N.A. | N.A. | *Fagus*<br>*sylvatica* | *Fagus*<br>*sylvatica* | *Ulex*<br>*europaeus* | *Festuca rubra* | *Festuca rubra* |
| Substrate type / Horizon | Organic peat<br>(Top layer) | Organic peat<br>(Top layer) | Organic soil | Gleic fluvisol | Mineral soil | Mineral soil | Litter horizon | Mineral soil (A<br>horizon) | Mineral soil | Mineral soil | Mineral soil |
| pH | 7.3 | 7.6 | 4.7 | 5.3 | 4.9 | 7.1 | N.A. | 4.5-5.1 | 3.9 | 5.6 | 5.9 |
| Moisture (% soil dry<br>weight) | 1010 | 900 | N.A | N.A. | 33 | 32 | 18 | 43-64 | 52 | 49 | 30 |
| # of replicates | 2 | 2 | 3 | 3 | 2 | 1 | 2 | 4 | 3 | 3 | 3 |
| Sampling time | August 2009 | August 2009 | July 2013 | July 2013 | July 2009 | January 2006 | May 2008 | May 2008 | March 2011 | March 2011 | March 2011 |
| Sequencing method | 454 GS FLX<br>Titanium | 454 GS FLX<br>Titanium | Illumina HiSeq<br>2500 | Illumina HiSeq<br>2500 | 454 GS FLX<br>Titanium | 454 GS 20 | 454 GS FLX | 454 GS FLX | Illumina HiSeq<br>2000 | Illumina HiSeq<br>2000 | Illumina HiSeq<br>2000 |

Furthermore, metatrascriptomic data were obtained from four different beech litter types (K, A, O and S), which had been incubated with the same microbial community in mesocosms (see Wanek et al., 2010 for details of the experimental setup). Litter samples were taken at three time points: after two weeks and after three and six months after inoculation, flash-frozen in liquid nitrogen and stored at −80 °C. RNA was extracted and double-stranded cDNA was prepared as previously described (Urich et al., 2008). 454 pyrosequencing was performed at the Norwegian Sequencing center, CEFS, University of Oslo (Norway). Raw sequence data were submitted to the NCBI Sequence Read Archive (SRA) under the accession number SRP134247.

### Bioinformatic analysis

Raw sequence datasets were filtered to a minimum length of 200 nucleotides and a minimum mean quality score of 25 using prinseq-lite (Schmieder & Edwards, 2011). SSU rRNA sequences were identified via SortMeRNA (Kopylova et al., 2012). USEARCH (Edgar, 2010) was used to randomly subsample datasets to a maximum of 50 000 - 100 000 sequences. The datasets were mapped against the CREST database silva123.1 by blastn (Altschul et al., 1990; Lanzén et al., 2012). The obtained blastn files were taxonomically analysed using MEGAN (Huson et al., 2011, min score 155; top percent 2.0; min support 1). The number of SSU rRNA reads of the investigated organisms was normalized in MEGAN to the total number of read counts. Investigated taxa with predatory lifestyle were *Myxococcales, Bdellovibrionales, Lysobacter, Dapterobacter, Vampirococcus, Amoebozoa, Cercozoa, Ciliophora, Foraminifera, Euglenozoa, Heterolobosea*, and *Nematoda*. Different Nematoda taxa were not investigated separately. The read counts of the analysed bacterial micropredators where substructed from the total bacterial SSU rRNA resulting in prey bacterial rRNA. The read counts of each analysed bacterivorous group were then normalized to the prey bacterial SSU rRNA reads.

Results for organic, excluding mofette (MO) samples, and mineral soils were tested for differentially expressed sequences with the R package edgeR (McCarthy et al., 2012; functions glmFit and glmLRT), using the non-normalized total read counts MEGAN file.

## Results

### Abundance of bacterivores in soil microbiomes

We screened the SSU (16S and 18S) rRNA fraction of 28 soil metatranscriptome datasets obtained from eleven different soils across Europe (Table 1) for bacterivorous pro- and eukaryotes. *Myxococcales* SSU rRNA reads comprised a high proportion of prey bacterial SSU rRNAs, ranging from 3.5 to 18.9% (9% on average), and higher than all other investigated bacterivores (Figure 1a). Their highest proportion in relation to bacteria was detected in peat soils. Additionally, an organic fluvisol and a beach litter layer showed *Myxococcales* abundances above 10%. The latter came up as the only exception in the pattern, i.e. here the Protozoa were the most abundant bacterivorous group. Overall, SSU rRNAs of protozoa were the second most abundant (Figure 1a). Like the *Myxococcales* they were generally more abundant in organic soils than in mineral soils. The only two cases where their proportion was above 10% of prey bacterial SSU rRNA reads were a peatland and a forest litter sample. While *Myxococcales* abundance never dropped below 3.4%, protozoa abundance was much lower in mineral soils (down to 0.7%). The third most abundant group were the *Nematoda* (Figure 1a). They showed greater variation in abundance compared to the aforementioned taxa, especially in organic soils, where they showed both their highest abundance (8.5%), namely in the forest litter horizon, and also their lowest abundance (< 0.1%), which occurred in the suboxic mofette soil. This was the only sampling site, where their abundance dropped below 0.1% of the prey bacterial SSU rRNA reads. The only other soils which showed *Nematoda* abundances above 1% were the organic peatland samples and the mineral Rothamsted soil. All mineral soils showed fractions of *Nematoda* SSU rRNAs within 0.1 – 1%. The *Bdellovibrionales* comprised even lower SSU rRNA abundances. Similar to the aforementioned, highest relative abundance of *Bdellovibrionales* was observed in organic soils (0.9% in peatland soil). We did not detect *Vampirococcus* and *Dapterobacter* in any of the investigated samples. *Lysobacter* comprised the lowest SSU rRNA abundances of all detected micropredators, namely 0.07% or lower.

**Figure 1.**
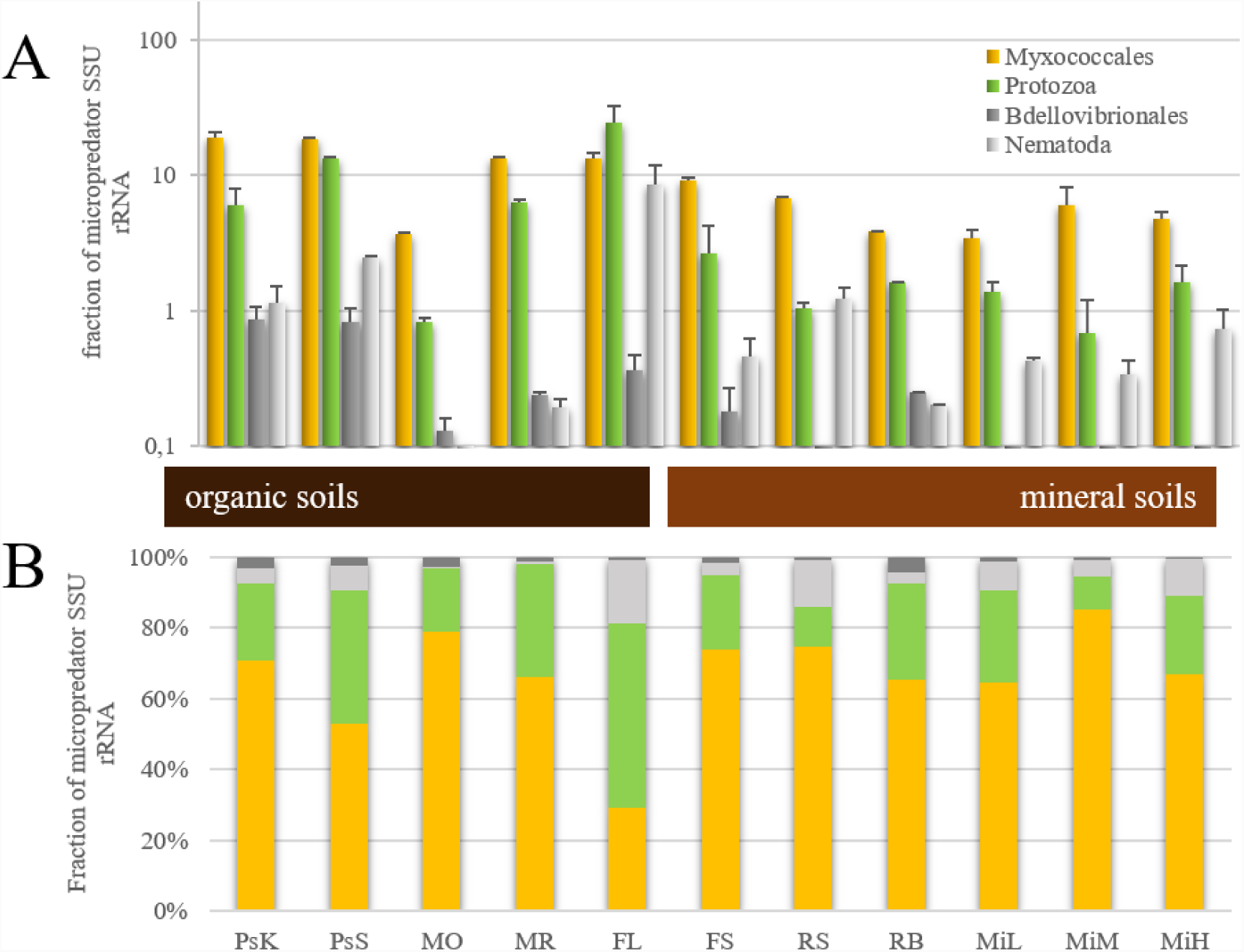
Screening of pro- and eukaryotic micropredators. (A) Fraction of major identified micropredator SSU rRNA normalized to SSU rRNA of prey bacteria. (B) Fraction of major identified micropredator SSU rRNA normalized to total micropredator SSU rRNA. Error bars show standard deviation of replicates. For sites see Table 1.

Looking at the composition of potential prey bacteria revealed that SSU rRNA from gram-negative bacteria comprised approximately 80%, while SSU rRNA from gram-positive were approximately 20%. The latter were slightly more abundant in mineral soils (supplementary figure S1).

### Myxococcales dominate bacterivorous taxa

Comparing all investigated bacterivorous groups, the *Myxococcales* were highest abundant in every sampling site, except for the forest litter (Figure 1b). In fact, in nine of the eleven sites, including all mineral sampling sites, the proportion of *Myxococcales* SSU rRNAs was more than 60% of all micropredators. In the forest litter their proportion of the bacterivorous groups was below 30%. Correspondingly, the protozoa were the most abundant group in that site, comprising up to more than half of all bacterial predators. However, in all the other sampled sites, the proportion of the protozoa was below 40%, in three cases even below 20%. Those were namely the organic mofette samples as well as the mineral Rothamsted site, and mine M from Spain, where the lowest percentage of all micropredators was observed. All of the sampling sites had *Nematoda* SSU rRNA below 20% Figure 1b). Their highest proportions occurred in the organic forest litter samples. Moreover, the only other two sites where their proportions were above 10%, were Rothamsted, where they were even more than the protozoa, and mineral mine H. All other sites showed proportions below 10%, with the lowest proportions in samples from mofette. The *Bdellovibrionales* were below 10% of micropredators in all sampling sites (Figure 1b).

### Community composition of Myxobacteria

We analysed the community composition of *Myxococcales* in more detail (Figure 2a). The most dominant family was *Haliangiaceae, followed by Polyangiaceae* and Blrii41, a family level group in the SILVA taxonomy that is currently devoid of cultured representatives. These three together comprised more than 2/3 of *Myxobacteria* SSU rRNAs in all but one site. *Haliangiaceae* and *Polyangiaceae* were more abundant in mineral soils, while Blrii41 was more characteristic for organic soils. The name-giving family *Myxococcaceae*, which is comprised, among others, of the most frequently isolated genera *Myxococcus* and *Corallococcus,* was barely present.

**Figure 2.**
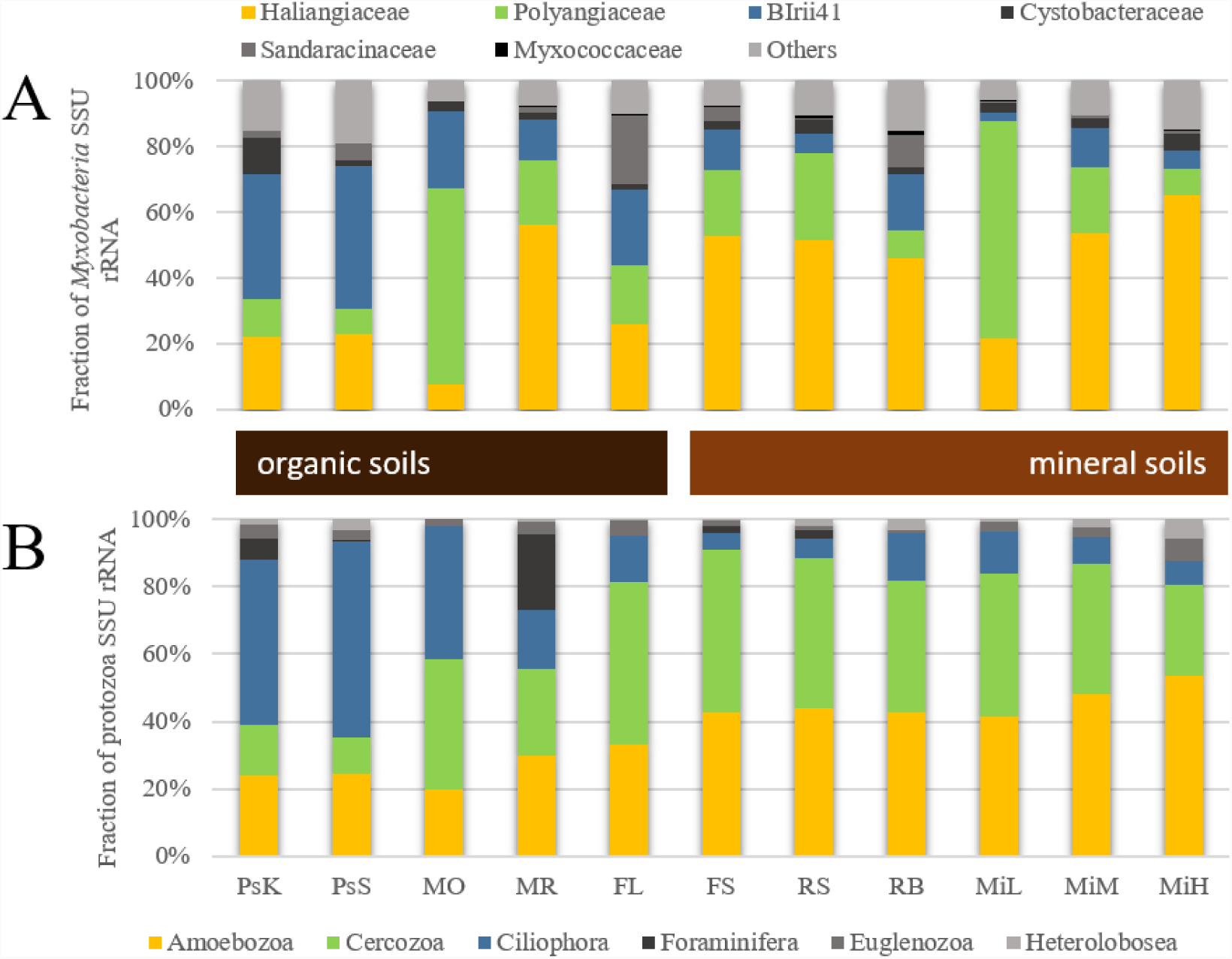
Screening of *Myxococcales* and protozoa taxa. (A) Fraction of identified *Myxococcales* SSU rRNA normalized to overall *Myxococcales* SSU rRNA. (B) Fraction of protozoa SSU rRNA normalized to total protozoa SSU rRNA. For sites see Table 1.

### Protozoa community composition

As previously found (Geisen et al., 2015), *Amoebozoa, Cercozoa* and *Ciliophora* were the three most abundant protist groups (Figure 2b). While *Amoebozoa* and *Cercozoa* dominated in mineral soils, the *Ciliophora* were most abundant in organic soils. The remaining predatory groups *Foraminifera, Euglenozoa*, and *Heterolobosea* accounted for low abundances on average. The mofette reference (MR) samples were an exception, with *Foraminifera* comprising more than 20% of protists.

### Dominance of Myxobacteria among bacterivores in mineral soils

We compared the average micropredator abundance (normalized to the prey bacteria) between mineral and organic soils (excluding mofette samples) based on SSU rRNA reads (Figure 3). *Lysobacter* data are not shown due to low abundances. Remarkably, micropredator SSU rRNAs comprised 32.3% of prey SSU rRNAs in organic soils, as compared to only 7.9% in mineral soils. Although concomitantly lower in abundance in mineral soils, *Myxococcales* comprised the highest micropredator proportions in both soil types (16.1% in organic vs. 5.7% in mineral soil). While protozoa were almost equally abundant in organic soil, they comprised approx. 1/4 of *Myxobacteria* in mineral soils. In fact, the percentage of *Myxococcales* within the bacterivores was remarkably higher in mineral soils, i.e. 72% compared to 50% in organic soils. Thus, the decrease in abundance of *Myxococcales* in mineral soils was not as strong as seen in the other micropredators.

**Figure 3.**
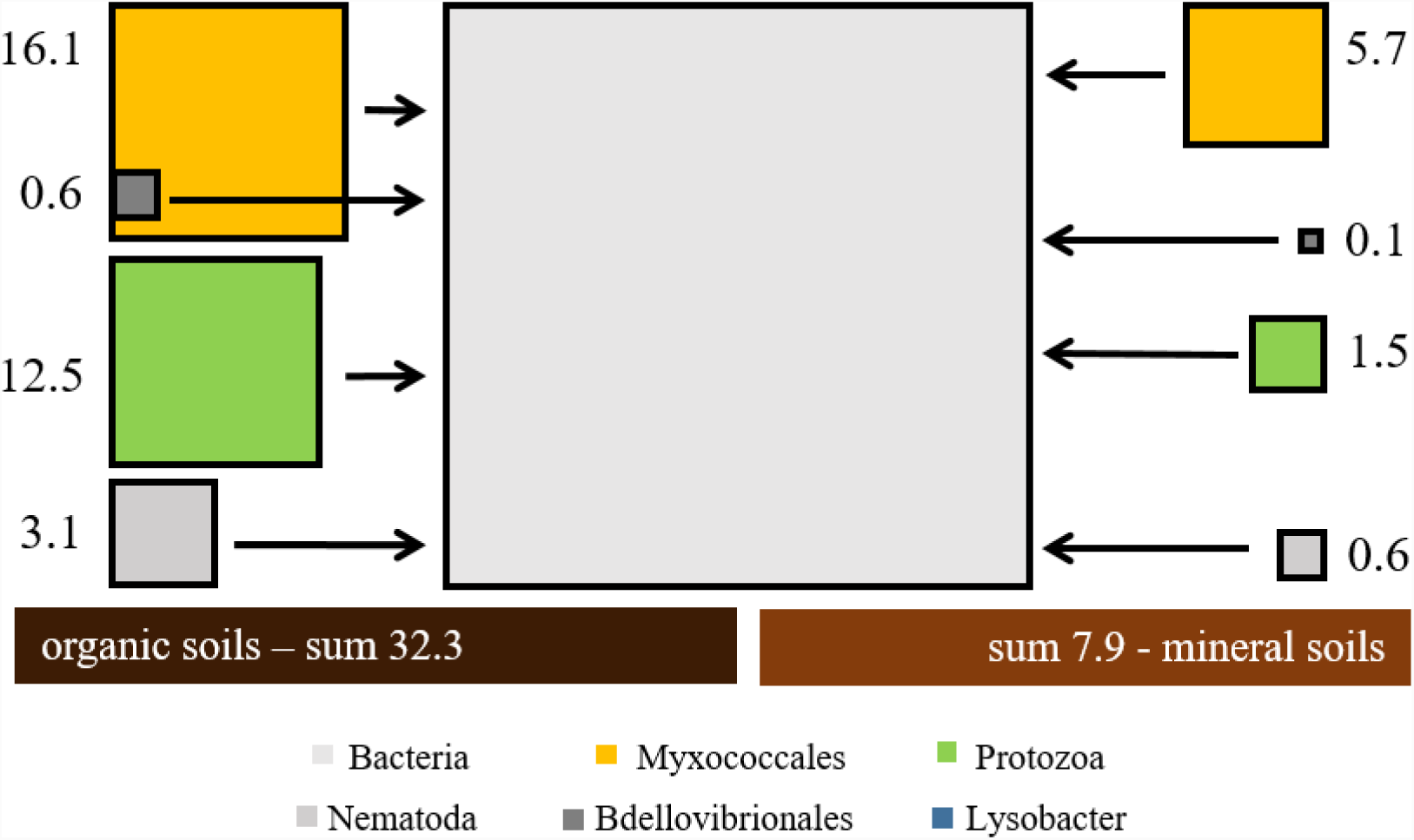
Comparison of organic and mineral soils. Fraction of major identified micropredator SSU rRNA normalized to SSU rRNA of prey bacteria. Average in organic soils (excluding MO samples) on the left; average in mineral soils in the right. Area of boxes resembles abundance of SSU rRNA. Numbers show proportions [%] of prey bacterial SSU rRNA. *Lysobacter* data are not shown due to low abundances.

To statistically verify the observed differences in abundance between organic and mineral soils, we tested the data for differentially expressed SSU rRNAs of micropredators. While the protozoa (p < 0.01), *Lysobacter* (p < 0.01), and *Bdellovibrionales* (p = 0.02) were significantly differently abundant between organic and mineral soils, no significant differences were detected for *Myxococcales* (p = 0.31) and *Nematoda* (p = 0.78). This supports the observed phenomenon, where the *Myxococcales* remained dominant in mineral soils, while the SSU rRNA abundances of other investigated micropredators significantly decreased in mineral soils.

### Temporal dynamics of bacterivores during community succession

It has been shown that *Myxobacteria* can have a saprotrophic life style next to bacterivory (reviewed in Reichenbach, 1999). We therefore analysed micropredator dynamics in a litter colonisation experiment. In a longitudinal experiment four types of sterilized beech litter differing in their C:N:P ratio were colonized by the same microbial community taken from beech forest soil (Supplementary Table 1 in Wanek et al., 2010). Metatranscriptome data were obtained from three time points over the course of six months. SSU rRNA abundances of protozoa, *Nematoda,* and *Myxococcales*, as well as total bacteria and fungi were assessed to investigate their temporal dynamics (Figure 4). Microbiomes on litters K and S, which had a higher nitrogen content, were strongly dominated by fungal SSU rRNAs as compared to bacterial rRNAs, while litters A and O, which had lower nitrogen content, had higher proportions of bacterial reads. The fungal:bacterial ratio stayed rather constant for each litter type over time. SSU rRNAs of bacterivores generally increased in relative abundance over time, especially from two weeks to three months. Remarkably, the bacterivores (including *Myxococcales*) appeared earlier and in higher relative abundance in litters with more prey bacteria. With few exceptions, protozoa comprised the most abundant predator of bacteria, with *Myxococcales* and *Nematoda* being second and third most abundant respectively. It appeared that after three months a rather stable predator:prey ratio had established.

**Figure 4.**
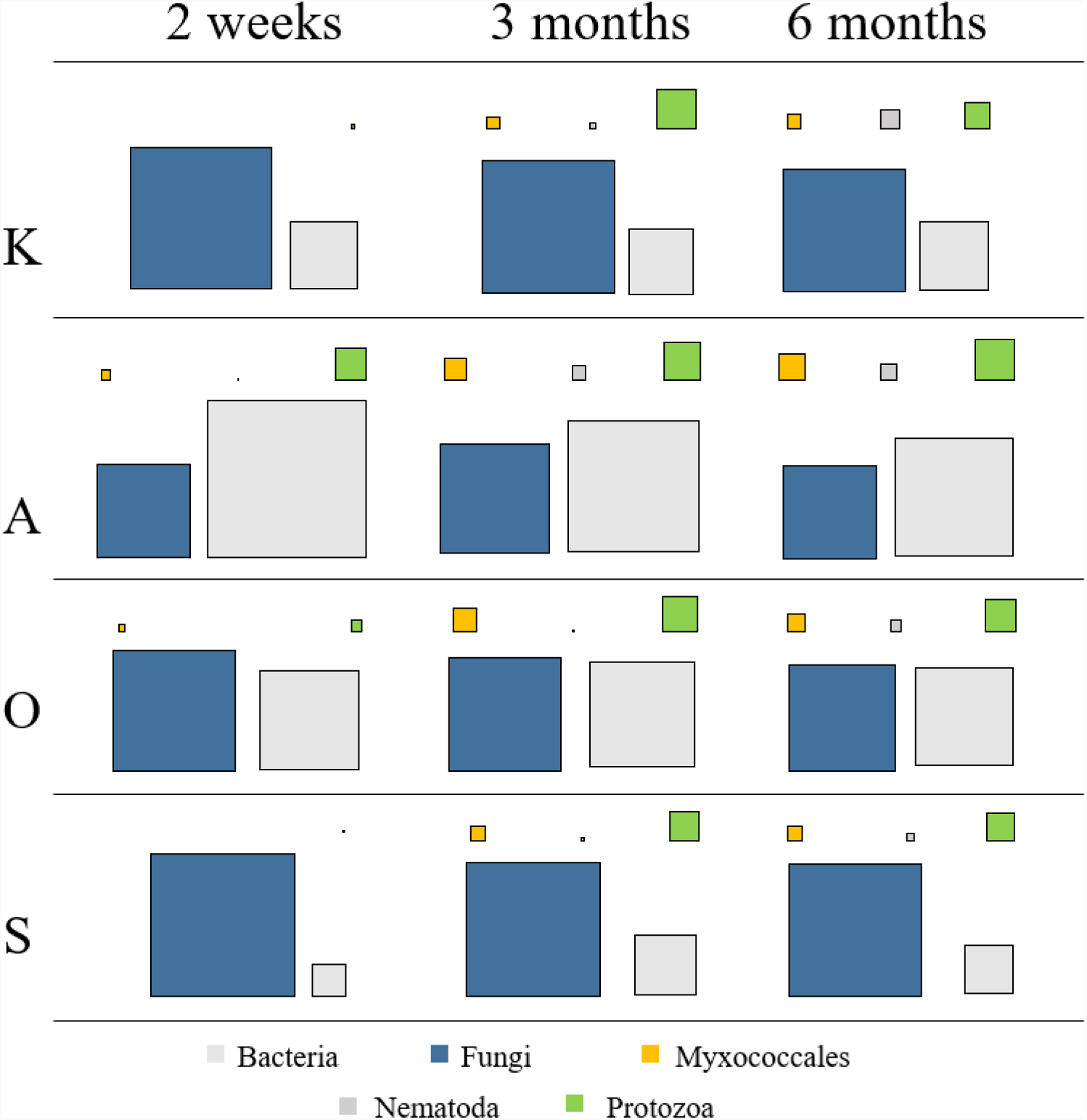
Colonisation of beech litter. Fraction of major identified micropredator, fungi, and prey bacteria SSU rRNA. Area of boxes resembles abundance of SSU rRNA. *Lysobacter* data are not shown due to low abundances. For litter types see Wanek *et al*., 2010.

## Discussion

### Metatranscriptomics-enabled holistic assessment of soil micropredators

It has been until recently impossible to assess soil bacterial and protist community composition with the same methodology. Although PCR amplicon approaches enabled the study of both groups separately, a direct comparison of their relative abundances was not possible due to the absence of universal primers that would tackle all groups without bias. The rRNA fraction of metatranscriptomics data enables broad three-domain community profiling of abundant bacteria, archaea, and eukaryotes via rRNA (Urich et al., 2008). This innovative approach has constantly been developed further and recently come to maturation due to lower-cost NGS sequencing technologies and bioinformatic tools (e.g. (Bengtsson et al., 2018; Schwab et al., 2014; Tveit et al., 2015)). Using this approach, we have recently created a first molecular census of active protists in soils (Geisen et al., 2015). The study showed that protozoa usually not detected with general PCR primers such as *Amoebozoa* and *Foraminifera* are abundantly present and provided the most comprehensive picture of active protist communities in soils to date. The strength of rRNA transcripts for comparatively unbiased views into community composition and assessment of unseen microbial diversity has recently gained popularity (e.g., Karst et al., 2018).

### Predatory Myxobacteria as key-stone taxon in mineral soils?

In all investigated soils the *Myxobacteria* comprised a significant proportion of the overall bacterial SSU rRNA transcripts. This confirms a recent PCR / 16S rRNA gene based survey where *Myxobacteria* also comprised a substantial fraction (4.1%; Zhou et al., 2014). In our study, lower micropredator abundances were detected in mineral soils as compared to organic soils, which may be due to less available carbon resulting in lower prey cell density in mineral soils. However, given the predatory traits of most myxobacterial taxa, their high relative abundance hints towards an important role in the microbial food web of soils. Moreover, with only one exception, the *Myxobacteria* exhibited the highest abundances when compared to all other bacterivores.

Traditionally, protists are considered to be the dominant group preying on bacteria (e.g. Geisen et al., 2016; Trap et al., 2016). In contrast to this, our data suggest an importance, possibly even dominance of *Myxococcales*. In fact, *Myxococcales* comprised approx. 3/4 of all micropredators in mineral soils. Possibly, the smaller pore sizes in mineral soils provide restricted access of protists to their bacterial prey, as compared to the smaller *Myxococcales*. Thus, microorganisms inhabiting non-continuous capillary pores could be protected from predation by Protozoa and *Nematoda*, but not from the similarly-sized *Myxobacteria*. The prokaryotes inhabiting the organic soil horizons, with unprotected macro-pore space, would in turn be subjected to higher grazing pressure. The *Myxobacteria* and protozoa exhibit fundamentally different predation strategies, with the much smaller *Myxobacteria* being famous for their social ‘wolf-pack’ hunting combined with the secretion of lytic enzymes, as compared to the larger phagotrophic protozoa (Reichenbach, 1999). The more similar cell size of *Myxobacteria* and prey bacteria could thus favour myxobacterial predation in mineral soils with small pores. Given the broad prey range of *Myxobacteria*, their high abundance in soils suggests a major influence on structuring the prokaryotic community composition, and might warrant their classification as key-stone taxon.

Interestingly, the majority of *Myxobacteria* was not related to the well-studied and easy-to-isolate *Myxococcaceae*, but belonged to families with a less well characterised prey spectrum, such as *Haliangiaceae, Kofleriaceae,* and *Polyangiaceae* (see Figure 2a), similar to findings of Zhou et al. (2014). The data in this study hint toward biases in culturability within the *Myxococcale*s. In fact, one large family-level group abundant in organic soils, represented in the SILVA database by clone Blrii41, is currently without any cultured representatives.

### A bacterial loop within the microbial loop

The soil survey could not give direct proof of whether the *Myxobacteria* (or any presumed micropredator) actually showed bacterivorous behavior *in situ*. In a colonisation experiment with sterilised beech litter we compared their succession with other bacterivores, such as the protozoa. In general, the abundance of potential bacterivores was positively associated with abundance of prey bacteria, and increased over time indicating a developing food web during litter colonisation. *Myxococcales* and protozoa abundances developed similarly over time. This observation hints at *Myxococcales* indeed having a predominantly predatory and not saprotrophic lifestyle, thus rendering the *Myxococcales* a prominent predatory taxon feeding on other bacteria.

There is direct *in situ* evidence for myxobacterial bacterivory from RNA-stable isotope probing studies. The hallmark study of Lueders and colleagues (2006) introduced the use of isotopically labelled prey bacteria to target the general diversity of micropredators in soil *in situ* and follow the carbon flow through the bacterial channel of the soil food web. The authors reported the detection of labelled sequences related to *Myxococcus, Lysobacter,* and *Bacteroidetes*. However, due to the limited technology at that time, they were not able to assess the contribution of predatory protozoa. Another shortcoming was the use of *E. coli* cells as prey because the survival of *E. coli* cells added to soil is rather low. In a more recent follow-up study with labelled native soil bacteria, Zhang and Lueders (2017) provided evidence for niche partitioning between bacterial and eukaryotic micropredators in soil, driven by the soil compartment. Interestingly, the *Myxobacteria* preyed on both gram-positive and gram-negative bacteria. Like in their previous study, the relative contribution of pro- and eukaryotic micropredators could not be assessed due to methodological limitations. Another recent study also included fungi and bacteria (Kramer et al., 2016), and indicated that the commonly accepted split of energy channels does not exist.

The dominance of *Myxobacteria*, especially in mineral soils, suggests their important role in the soil microbial food web (Figure 5). The so-called microbial loop in soil (Bonkowski, 2004) is important for the remineralisation of nutrients, where especially protozoa and *Nematoda* feed on bacteria and by this set free nutrients, which are in turn provided for the bacteria as well as plants (Coleman, 1994). Our observations hint at the presence of an ecologically important ‘bacterial loop’, especially in mineral soils, within the prokaryotes and independent of protozoa, that has been overlooked until today (Figure 5). As a consequence, bacterial micropredators might not only be important for shaping microbial communities but might also prove to be important for the recycling of nutrients in soils, as it has been shown for protozoa (Bonkowski, 2004; Koller et al., 2013), and thus potentially for nutrient and carbon cycling.

**Figure 5.**
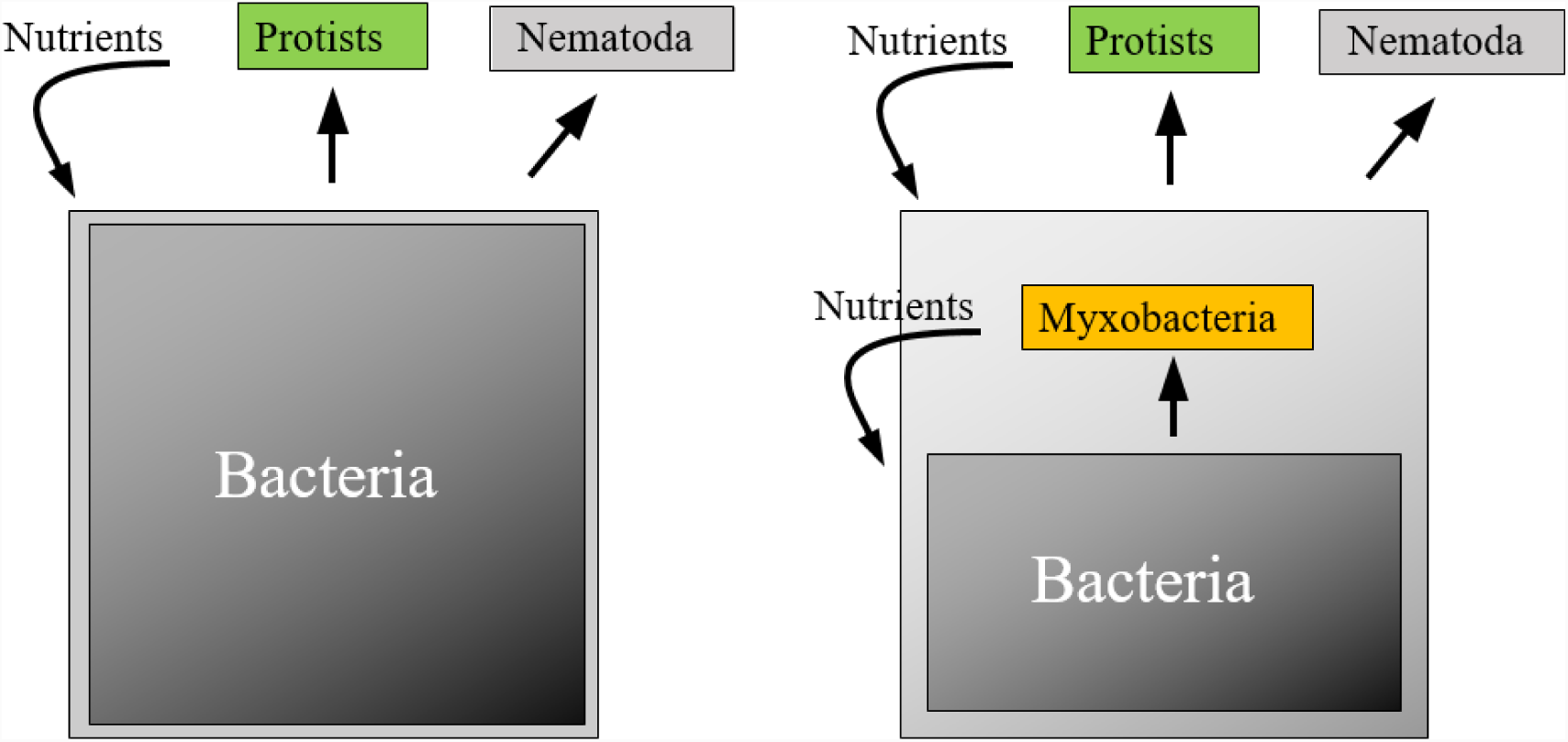
Simplified soil microbial loop. Left: Traditional microbial loop with separate roles of prokaryotic and eukaryotic organisms. Right: Microbial loop containing a bacterial loop independent of eukaryotic organisms. Straight arrows: links between trophic levels. Bent arrows: provision of nutrients.

Although the *Myxococcales* order comprised a significantly high proportion of bacterial SSU rRNA in all sites, differences at family-level were observed among the soils. Nevertheless, the two most abundant families were the *Polyangiaceae* and *Haliangiaceae*. The high dominance of the latter may well be due to the *Haliangiaceae* taxon being clustered together with the *Kofleriaceae* taxon in the current SILVA database. Interestingly the name-giving group, the *Myxococcaceae,* comprised a very low proportion of all *Myxococcales*. The *Myxococcaceae* are known to be easily cultivable from a variety of environmental samples. Our data show that other, less well characterised families are in fact much more abundant in soil. Thus, efforts should be undertaken to investigate their biology and in particular their prey spectrum.

### Methodological considerations

The micropredator abundance data in this study are derived from the abundance of SSU rRNA in metatranscriptomes. This does not reflect organismic abundance but is rather a proxy of living biomass (Urich & Schleper, 2011). In fact, several factor need to be taken into account when comparing the SSU rRNA from different pro- and eukaryotic organisms. Results of various studies suggest differences in RNA contents per biomass (1) between organisms and (2) between growth phases, respectively. The ribosome density in prokaryotic cells is generally considered higher than in eukaryotes. However, few data are available to our knowledge. The RNA content of *E. coli* was determined to be 15.7% of dry mass (dm) (Stouthamer & van Leeuwenhoek, 39,545-565, 1973), of *Bacillus subtilis* between 8.5% and 14% dm^-1^ (Tempest et al., 1968), *Saccaromyces cerevisiae* 23% dm^-1^ (Parada & Acevedo, 1983), *Aspergillus* 5.9% (Carlsen et al., 2000) and *Penicillium chrysogenum* between 5% and 8% (Henriksen et al., 1996). Furthermore, prokaryotic cells in an exponential growth phase are known to contain more RNA than cells in stationary phase (e.g. Tempest et al., 1968). Preliminary data (Petters & Urich, unpublished) hint to correction factors to be applied when comparing rRNA based abundances from metatranscriptomes between pro- and eukaroytes. Nevertheless, a recent study showed that rRNA correlates better with cell counts than ribosomal RNA genes (Giner et al., 2016). Thus, our metatranscriptomics data identify the predatory *Myxobacteria* as important players in the midst of the soil food web and suggests a prominent role in the soil microbial loop in particular.

## Acknowledgements

Andreas Richter, Wolfgang Wanek and colleagues (University of Vienna) are thanked for providing beech litter samples. Daniela Teichmann and Sylvia Klaubauf (University of Vienna) are acknowledged for excellent technical assistance in RNA extraction. We thank Ave Tooming-Klunderud, Lex Nederbragt and others at the Norwegian High-Throughput Sequencing Centre (University of Oslo) for 454 pyrosequencing. Andrea Söllinger acknoledges funding from the University of Vienna (uni:docs) and the OeAD (Austrian agency for international mobility and cooperation in education, science and research; Marietta-Blau-Fellowship).

## Authors’ contributions

The study was designed by TU. Data analysis was performed by SP and TU, supported by AS and MB. The manuscript was written by SP and TU, assisted by all co-authors.

